## Supplemental Figures and Text for "Allosteric activation of CwlD amidase activity by the GerS lipoprotein during *Clostridioides difficile* spore formation"

### Supplementary Material Description

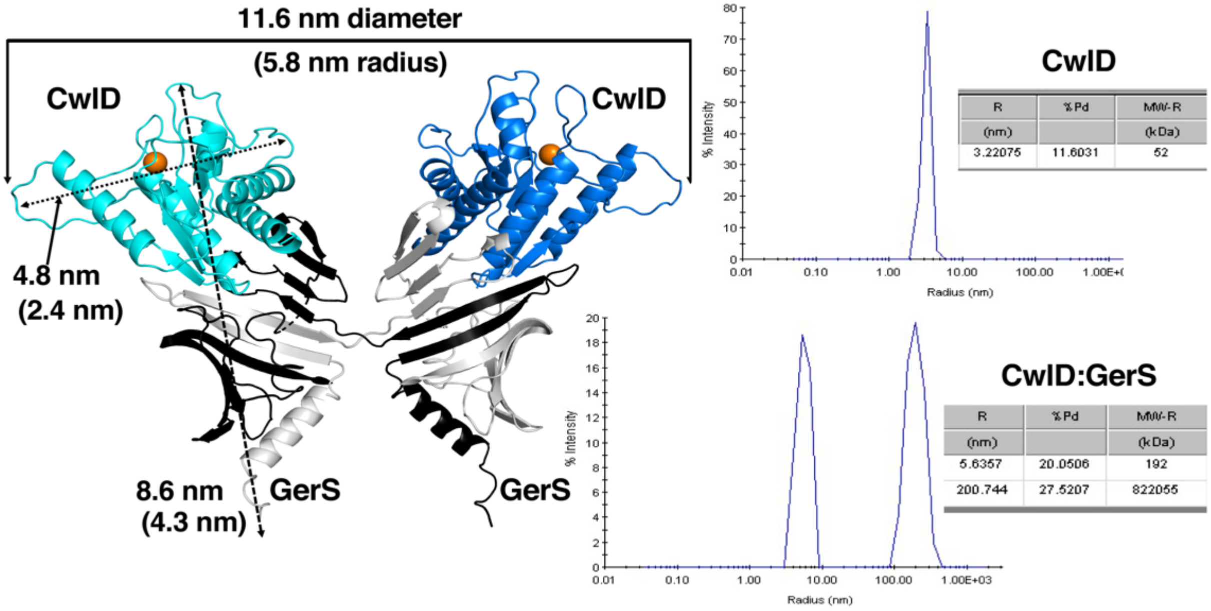


**Figure S1. Comparison of the CwlD:GerS complex to solution behavior.** The left panel shows the two CwlD / two GerS complex within the crystal with dimensions along each axis shown in nm. Dynamic light scattering results are shown for the purified CwlD and the CwlD:GerS complex. The radius of the CwlD:GerS complex in solution of 5.6 nm agrees well with 5.8 nm as measured in the crystal. The 200 nm peak in the complex sample is excess GerS not in the complex as GerS in the absence of CwlD behaves multimodally with some aggregation.

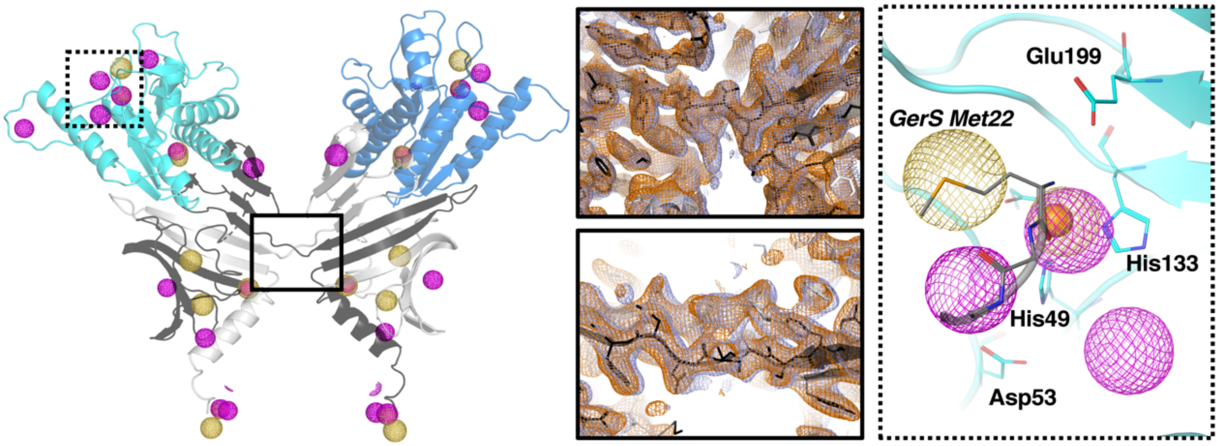


**Figure S2. Heavy atom phasing of the CwlD:GerS complex structure.** Left shows the two CwlD (Blue tones) / 2 GerS (grey tones) complex . The yellow mesh shows the anomalous signal at 3 sigma for the selenomethionine derivative and the maroon the sodium iodide derivative from Autosharp (Global Phasing Limited). Middle panels show Se-SAD phases in blue and Se/NaI with native MIR phases in orange contoured at 1 sigma and the beta strand crossover between the two GerS protomers (solid box) with views 90$^{o}$ apart. Right panel (dotted box) shows the anomalous peaks near the zinc binding site of CwlD.

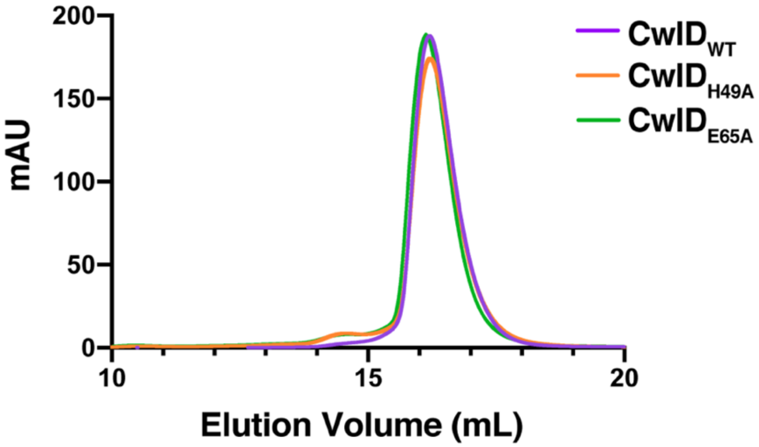


**Figure S3. Size Exclusion Chromatography analyses of CwlD variants.** Purified His-tagged CwlD$_{WT}$, CwlD$_{H49A}$, and CwlD$_{E65A}$ were analyzed by size exclusion chromatography. mAU corresponds to the UV absorbance measurements ($A_{280}$) during the protein elution.

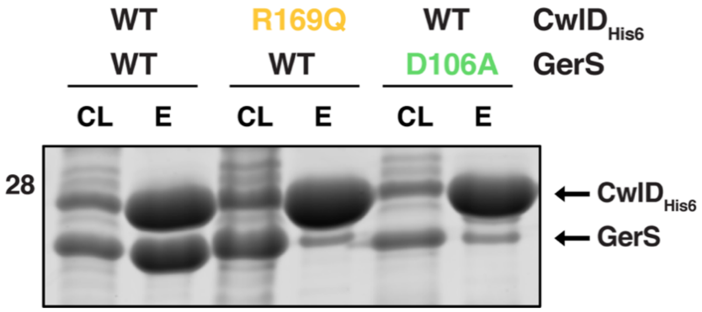


**Figure S4. Coomassie stain of co-affinity purifications of His-tagged CwlD$_{WT}$ or CwlD$_{R169Q}$ with either GerS$_{WT}$ or GerS$_{D106A}$.** The indicated proteins were produced in *E. coli* and purified using $Ni^{2+}$- affinity resin. Cleared lysate (CL) and eluate (E) fractions were analyzed using Coomassie staining.

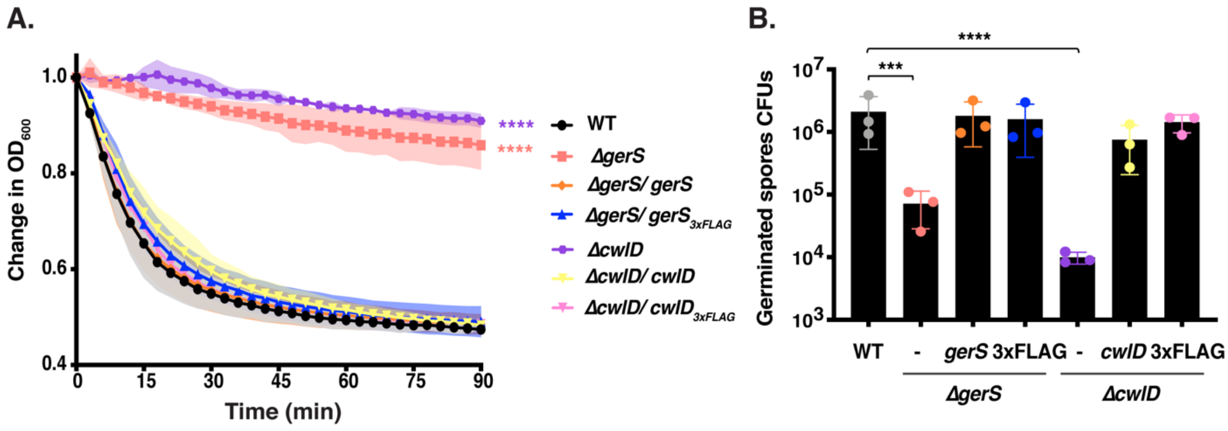


**Figure S5. Germination efficiency of CwlD$_{\mathrm{FLAG}}$ and GerS$_{\mathrm{FLAG}}$ complementation strains.** (A) Change in the $OD_{600}$ in response to germinant of Δ*gerS* and Δ*cwlD* spores complemented with *gerS*$_{\mathrm{FLAG}}$ and *cwlD*$_{\mathrm{FLAG}}$ constructs, respectively, relative to spores complementated with native *gerS* or *cwlD*, respectively. Purified spores were resuspended in BHIS, and germination was induced by adding taurocholate (1% final concentration). The ratio of the $OD_{600}$ of each strain at a given time point relative to the $OD_{600}$ at time zero is plotted. The mean of three assays from 3 independent spore preparations are shown. Shading represents the standard deviation as the area between error bars for each time point measured. Statistical significance relative to wild-type was determined using two-way ANOVA and Tukey's test. **** $p < 0.0001$. (B) Spore germination efficiency of the strains relative complemented with FLAG-tagged vs. wild-type complementation constructs. The number of colony forming units (CFUs) produced by germinating spores is shown. Average of results from three independent spore preparations are shown along with the associated standard deviations. Statistical analyses relative to the wild-type were performed using one-way ANOVA and Tukey's test. ***, $p < 0.001$; **** $p < 0.0001$.

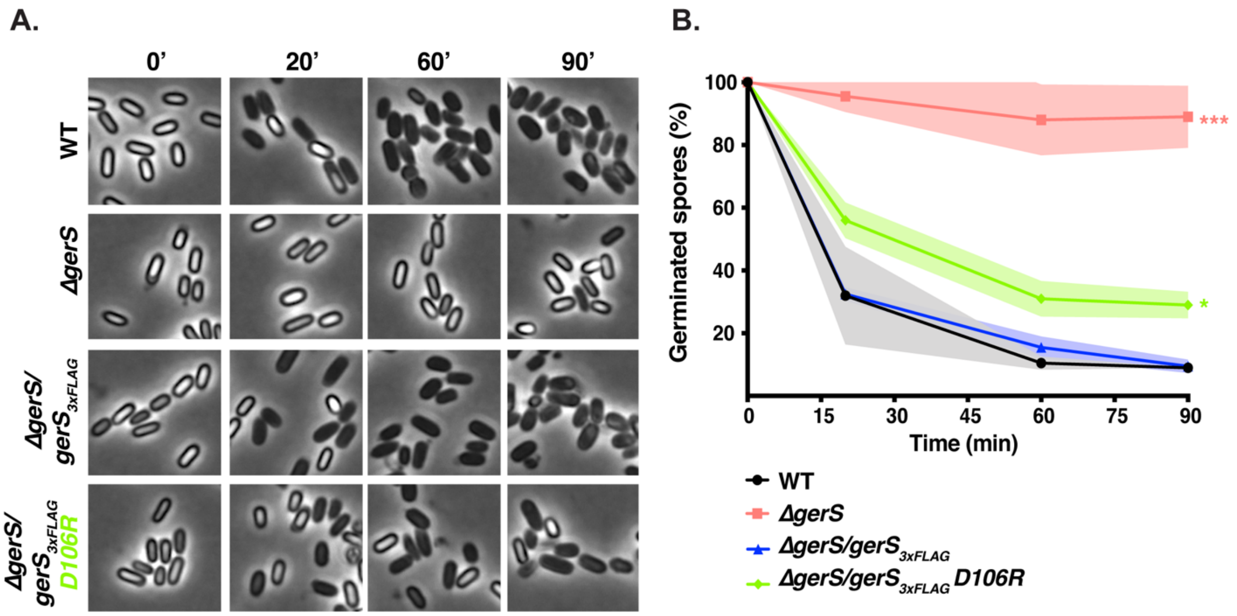


**Figure S6. Spore germination of strains encoding FLAG-tagged GerS$_{D106R}$ variants monitored by phase-contrast microscopy.** (A) Purified spores were resuspended in BHIS ± germinant and incubated aerobically for 90 min at 37°C. At the indicated time points samples were fixed in paraformaldehyde and visualized using phase-contrast microscopy. Scale bar, 1 um. B) Percent spore germination over time as detected by phase-contrast microscopy. The percentage of phase-bright spores of each strain at a given time point relative to the percentage at time zero is plotted. The mean of three assays from 2 independent spore preparations are shown. Shading represents the standard deviation as the area between error bars for each time point measured. Statistical significance relative to wild-type was determined using two-way ANOVA and Tukey's test. **** $p < 0.0001$.

### Supplementary Text S1 – *E. coli* strain construction

**pET22b-*cwlD*$_{\Delta 25}$-H49A-His$_6$.** Primer pair #2545 and 2763 and primer pair #2762 and 2546 were used to amplify regions spanning the *cwlD* gene lacking its stop codon and first 25 codons. Primers 2763 and 2762 encode the H49A mutation. The PCR products were cloned into pET22b digested with NdeI/SalI using Gibson assembly. The Gibson was transformed into DH5α, and the resulting construct was sequenced verified before transforming the plasmid into BL21(DE3) for protein expression.

**pET22b-*cwlD*$_{\Delta 25}$-E65A-His$_6$.** Primer pair #2545 and 3309 and primer pair #3310 and 2546 were used to amplify regions spanning the *cwlD* gene lacking its stop codon and first 25 codons. Primers 3309 and 3310 encode the E65A mutation. The PCR products were cloned into pET22b digested with NdeI/SalI using Gibson assembly. The Gibson was transformed into DH5α, and the resulting construct was sequenced verified before transforming the plasmid into BL21(DE3) for protein expression.

**pET22b-*cwlD*$_{\Delta 25}$-E199A-His$_6$.** Primer pair #2545 and 3251 and primer pair #3252 and 2546 were used to amplify regions spanning the *cwlD* gene lacking its stop codon and first 25 codons. Primers 3251 and 3252 encode the E199A mutation. The PCR products were cloned into pET22b digested with NdeI/SalI using Gibson assembly. The Gibson was transformed into DH5α, and the resulting construct was sequenced verified before transforming the plasmid into BL21(DE3) for protein expression.

**pET22b-*cwlD*$_{\Delta 25}$-R169D-His$_6$.** Primer pair #2545 and 3731 and primer pair #3732 and 2546 were used to amplify regions spanning the *cwlD* gene lacking its stop codon and first 25 codons. Primers 3731 and 3732 encode the R169D mutation. The PCR products were cloned into pET22b digested with NdeI/SalI using Gibson assembly. The Gibson was transformed into DH5α, and the resulting construct was sequenced verified before transforming the plasmid into BL21(DE3) for protein expression.

**pET22b-*cwlD*$_{\Delta 25}$-R169Q-His$_6$.** Primer pair #2545 and 3727 and primer pair #3728 and 2546 were used to amplify regions spanning the *cwlD* gene lacking its stop codon and first 25 codons. Primers 3727 and 3728 encode the R169Q mutation. The PCR products were cloned into pET22b digested with NdeI/SalI using Gibson assembly. The Gibson was transformed into DH5α, and the resulting construct was sequenced verified before transforming the plasmid into BL21(DE3) for protein expression.

**pET22b-*cwlD*$_{\Delta 25}$-E78Q-His$_6$.** Primer pair #2545 and 3733 and primer pair #3734 and 2546 were used to amplify regions spanning the *cwlD* gene lacking its stop codon and first 25 codons. Primers 3733 and 3734 encode the E78Q mutation. The PCR products were cloned into pET22b digested with NdeI/SalI using Gibson assembly. The Gibson was transformed into DH5α, and the resulting construct was sequenced verified before transforming the plasmid into BL21(DE3) for protein expression.

**pET22b-*gerS*$_{\Delta 22}$-His$_6$.** Primer pair #3103 and 3104 was used to amplify the *gerS* gene lacking its stop codon and first 22 codons. The PCR product was cloned into pET22b digested with NdeI/SalI using Gibson assembly. The Gibson was transformed into DH5α, and the resulting construct was sequenced verified before transforming the plasmid into BL21(DE3) for protein expression.

**pET29a-*gerS*$_{\Delta 22}$-H61A.** Primer pair #3419 and 3725 and primer pair #3726 and 3420 were used to amplify regions spanning the *gerS* gene lacking its first 22 codons. Primers 3725 and 3726 encode the H61A mutation. The PCR products were cloned into pET29a digested with NdeI/XhoI using Gibson assembly. The Gibson was transformed into DH5α, and the resulting construct was sequenced verified before transforming the plasmid into BL21(DE3) for protein expression.

**pET29a-*gerS*$_{\Delta 22}$-D106R.** Primer pair #3419 and 3723 and primer pair #3724 and 3420 were used to amplify regions spanning the *gerS* gene lacking its first 22 codons. Primers 3723 and 3724 encode the D106R mutation. The PCR products were cloned into pET29a digested with NdeI/XhoI using Gibson assembly. The Gibson was transformed into DH5α, and the resulting construct was sequenced verified before transforming the plasmid into BL21(DE3) for protein expression.

**pET29a-*gerS*$_{\Delta 22}$-D106N.** Primer pair #3419 and 3719 and primer pair #3720 and 3420 were used to amplify regions spanning the *gerS* gene lacking its first 22 codons. Primers 3719 and 3720 encode the D106N mutation. The PCR products were cloned into pET29a digested with NdeI/XhoI using Gibson assembly. The Gibson was transformed into DH5α, and the resulting construct was sequenced verified before transforming the plasmid into BL21(DE3) for protein expression.

**pMTL-YN1C-*cwlD*$_{H49A}$.** Primer pair #2362 and 2763 and primer pair #2762 and 2450 were used to amplify regions spanning 274 bp upstream of *cwlD* and the *cwlD* gene including the stop codon.
Primers 2763 and 2762 encode the H49A mutation. The PCR products were cloned into pMTL-YN1C digested with NotI/XhoI using Gibson assembly.

**pMTL-YN1C-*cwlD*$_{E65A}$.** Primer pair #2362 and 3309 and primer pair #3310 and 2450 were used to amplify regions spanning 274 bp upstream of *cwlD* and the *cwlD* gene including the stop codon.
Primers 3309 and 3310 encode the E65A mutation. The PCR products were cloned into pMTL-YN1C digested with NotI/XhoI using Gibson assembly.

**pMTL-YN1C-*cwlD*$_{E199A}$.** Primer pair #2362 and 3251 and primer pair #3252 and 2450 were used to amplify regions spanning 274 bp upstream of *cwlD* and the *cwlD* gene including the stop codon.
Primers 3251 and 3252 encode the E199A mutation. The PCR products were cloned into pMTL-YN1C digested with NotI/XhoI using Gibson assembly.

**pMTL-YN1C-*cwlD*-3xFLAG**. To clone the *cwlD* complementation construct encoding a C-terminal $FLAG_3$ epitope tag, primer pair #2362 and 2599 was used to amplify the *cwlD* gene without the stop codon and its promoter region (274 bp upstream) along with sequence encoding part of a FLAG epitope. The resulting PCR product was assembled with the following g-block encoding the $FLAG_3$ epitope into pMTL-YN1C along with digested with NotI and XhoI using Gibson assembly.

gBlock: Cdif CwlD-3xFLAG for pMTL-YN1C
CAGAGAAGTCAAACCAAGGGATGATATATATCTTTTGAAAGACAATAATATTCCATCAGTAC
TGATAGAATGTGGTTTTTATCAAATGAAAAAGAGTGTAAACTCTTAACTGATGAAACATAT
CAAGAAAAAATAGCATGGGCAATCTACATAGGAATACAAAAATATTTAAGTGATTATAAAG
ATGATGATGATAAAGACTATAAAGATGACGATGATAAGGATTATAAGGATGATGATGACAA
ATAACTCGAGGCCTGCAGACATGCAAGC

**pMTL-YN1C-*cwlD*$_{H49A}$-3xFLAG**. Primer pair #2362 and 2763 and primer pair #2762 and 2562 were used to amplify the *cwlD* gene without the stop codon and its promoter region (274 bp upstream) along with sequence encoding the $FLAG_3$ epitope using the pMTL-YN1C-*cwlD*-3xFLAG as template. Primers 2763 and 2762 encode the H49A mutation. The PCR products were cloned into pMTL-YN1C digested with NotI/XhoI using Gibson assembly.

**pMTL-YN1C-*cwlD*$_{E199A}$-3xFLAG**. Primer pair #2362 and 3251 and primer pair #3252 and 2562 were used to amplify the *cwlD* gene without the stop codon and its promoter region (274 bp upstream) along with sequence encoding the $FLAG_3$ epitope using the pMTL-YN1C-*cwlD*-3xFLAG as template. Primers 3251 and 3252 encode the E199A mutation. The PCR products were cloned into pMTL-YN1C digested with NotI/XhoI using Gibson assembly.

**pMTL-YN1C-*cwlD*$_{R169D}$-3xFLAG.** Primer pair #2362 and 3731 and primer pair #3732 and 2562 were used to amplify the *cwlD* gene without the stop codon and its promoter region (274 bp upstream) along with sequence encoding the $FLAG_3$ epitope using the pMTL-YN1C-*cwlD*-3xFLAG as template. Primers 3731 and 3732 encode the R169D mutation. The PCR products were cloned into pMTL-YN1C digested with NotI/XhoI using Gibson assembly.

**pMTL-YN1C-*gerS*-3xFLAG-*alr2*.** To clone the *gerS* complementation construct encoding a C-terminal $FLAG_3$ epitope tag, primer pair #2181 and 2365 was used to amplify the *gerS* gene without the stop codon and its promoter region (367 bp upstream) along with sequence encoding part of a FLAG epitope. The resulting PCR product was assembled with the following g-block encoding the $FLAG_3$ epitope. The *alr2* gene was amplified using primer pair #2617 and 2408. The two PCR fragments were cloned into pMTL-YN1C digested with NotI and XhoI using Gibson assembly.

gBlock: Cdif GerS-3xFLAG-alr2 for pMTL-YN1C

GAGGAGATGAAAGTAAAGTTATCCAAAGGACATTTGGTTTTAGAGACTTTTATTCCTGGGGA
TAATAAATATTTTAACAAGCAAGTATTATATGTAAATGCTGACACAAAAAATCCTGAAAAAA
TGGAAGTGTTAGATAAAGAGGGAGTGCCAAGATTTACAGTAAAATACAAAGATTTTGAATA
CAGAAACGATTATAAGGATGACGATGATAAAGACTATAAAGATGACGATGATAAGGATTAT
AAGGATGACGATGACAAATAACTCGAGGCCTGCAGACATGCAAGCTTGGC

**pMTL-YN1C-*gerS*$_{D106R}$-3xFLAG-*alr2*.** Primer pair #2181 and 3723 and primer pair #3724 and 2408 were used to amplify the *gerS* gene without the stop codon and its promoter region (367 bp upstream) along with sequence encoding the $FLAG_3$ epitope and the *alr2* gene using the pMTL-YN1C-*cwlD*-3xFLAG-*alr2* as template. Primers 3723 and 3724 encode the D106R mutation. The PCR products were cloned into pMTL-YN1C digested with NotI/XhoI using Gibson assembly.

**Supplementary Table S1. Strains and plasmids used in this study.**

| Strain # | Strain name | Relevant genotype or features | Source/ reference |
|---|---|---|---|
| ***C. difficile* strains – 630Δerm** | | | |
| 756 | 630Δ*erm*Δ*pyrE* | *erm*-sensitive derivate of 630 with a deletion in *pyrE* | (1) |
| 846 | 630Δ*erm*-p | *erm*-sensitive derivate of 630 with *pyrE* restored | (2) |
| 849 | 630Δ*erm* Δ*spo0A*-p | 630Δ*erm* Δ*spo0A* with *pyrE* restored | (2) |
| 950 | 630Δ*erm* Δ*gerS*Δ*pyrE* | 630Δ*erm*Δ*pyrE* with *gerS* deleted (*CD630_34640*) | (3) |
| 1075 | 630Δ*erm* Δ*gerS*-p | 630Δ*erm* Δ*gerS* with *pyrE* restored | (3) |
| 1617 | 630Δ*erm* Δ*cwlD*Δ*pyrE* | 630Δ*erm*Δ*pyrE* with *cwlD* deleted (*CD630_01060*) | (3) |
| 1677 | 630Δ*erm* Δ*gerS*/*gerS-alr2* | 630Δ*erm* Δ*gerS* with *gerS-alr2* in the *pyrE* locus | (3) |
| 1686 | 630Δ*erm* Δ*cwlD*-p | 630Δ*erm* Δ*cwlD* with *pyrE* restored | (3) |
| 1692 | 630Δ*erm* Δ*cwlD*/*cwlD* | 630Δ*erm* Δ*cwlD* with *cwlD* in the *pyrE* locus | (3) |
| 2053 | 630Δ*erm*- *p*-ΔgerS/*gerS*-3xFLAG-*alr2* | 630Δ*erm* ΔgerS with gerS-3xFLAG-*alr2* in the *pyrE* locus | This study |
| 2056 | 630Δ*erm* Δ*cwlD*/*cwlD*-3xFLAG | 630Δ*erm* Δ*cwlD* with *cwlD*-3xFLAG in the *pyrE* locus | This study |
| 2161 | 630Δ*erm* Δ*cwlD*/$cwlD_{H49A}$ | 630Δ*erm* Δ*cwlD* with $cwlD_{H49A}$ in the *pyrE* locus | This study |
| 2764 | 630Δ*erm* Δ*cwlD*/$cwlD_{E199A}$ | 630Δ*erm* Δ*cwlD* with $cwlD_{E199A}$ in the *pyrE* locus | This study |
| 2900 | 630Δ*erm* Δ*cwlD*/$cwlD_{E65A}$ | 630Δ*erm* Δ*cwlD* with $cwlD_{E65A}$ in the *pyrE* locus | This study |
| 2918 | 630Δ*erm* Δ*cwlD*/$cwlD_{H49A}$-3xFLAG | 630Δ*erm* Δ*cwlD* with $cwlD_{H49A}$-3xFLAG in the *pyrE* locus | This study |
| 2924 | 630Δ*erm* Δ*cwlD*/$cwlD_{E199A}$-3xFLAG | 630Δ*erm* Δ*cwlD* with $cwlD_{E199A}$-3xFLAG in the *pyrE* locus | This study |
| 3549 | 630Δ*erm* Δ*gerS*/$gerS_{D106R}$-3xFLAG | 630Δ*erm* Δ*cwlD* with $gerS_{D106R}$-3xFLAG-*alr2* in the *pyrE* locus | This study |
| 3564 | 630Δ*erm* Δ*cwlD*/$cwlD_{R169D}$-3xFLAG | 630Δ*erm* Δ*cwlD* with $cwlD_{R169D}$-3xFLAG in the *pyrE* locus | This study |
| ***E. coli* strains** | | | |
| 41 | DH5a | F– Φ80*lacZ*ΔM15 Δ(*lacZYA-argF*) U169 *recA1 endA1 hsdR17* ($rK^-$, $mK^+$) *phoA supE44* λ– *thi-1 gyrA96 relA1* | D. Cameron |
| 531 | HB101/pRK24 | F- *mcrB mrr hsdS20*($rB^-mB^-$) *recA13 leuB6 ara-13 proA2 lavYI galK2 xyl-6 mtl-1 rpsL20* carrying pRK24 | C. Ellermeier |
| 892 | BL21 (DE3) | $F^-$ *ompT hsdSB*($rB^-mB^-$) *gal dcm* (DE3) | C. Ellermeier |
| | B834 (DE3) | $F^-$ *ompT* $hsdS_B$($r_B^-$ $m_B^-$) *gal dcm met* (DE3) | Novagen |
| 2045 | pET22b-Δ25-*cwlD*-$His_6$ | pET22b-*cwlD* (deletion of N-terminal 25 aa with His6 tag) in BL21(DE3) | (3) |
| 2147 | pMTL-YN1C-*gerS*-3xFLAG-*alr2* | pMTL-YN1C- *gerS*-3xFLAG in HB101/pRK24 | This study |
| 2149 | pMTL-YN1C-*cwlD*-3xFLAG | pMTL-YN1C-*cwlD*-3xFLAG in HB101/pRK24 | This study |
| 2191 | pMTL-YN1C-$cwlD_{H49A}$ | pMTL-YN1C-$cwlD_{H49A}$ in HB101/pRK24 | This study |
| 2219 | pET22b-Δ25-$cwlD_{H49A}$-$His_6$ | pET22b-*cwlD* (deletion of N-terminal 25 aa and H49A point mutation with His6 tag) in BL21(DE3) | This study |
| 2395 | pET22b-Δ22-*gerS*-$His_6$ | pET22b-gerS (deletion of N-terminal 22 aa with His6 tag) in BL21(DE3) | This study |
| 2493 | pMTL-YN1C-$cwlD_{E199A}$ | pMTL-YN1C-$cwlD_{E199A}$ in HB101/pRK24 | This study |

| | | | |
|---|---|---|---|
| 2557 | pMTL-YN1C-*cwlD*$_{E65A}$ | pMTL-YN1C-*cwlD*$_{E65A}$ in HB101/pRK24 | This study |
| 2566 | pMTL-YN1C-*cwlD*$_{H49A}$-3xFLAG | pMTL-YN1C-*cwlD*$_{H49A}$-3xFLAG in HB101/pRK24 | This study |
| 2570 | pMTL-YN1C-*cwlD*$_{E199A}$-3xFLAG | pMTL-YN1C-*cwlD*$_{E199A}$-3xFLAG in HB101/pRK24 | This study |
| 2574 | pET22b-Δ25-*cwlD*$_{E65A}$-$His_6$ | pET22b-*cwlD* (deletion of N-terminal 25 aa and E65A point mutation with His6 tag) in BL21(DE3) | This study |
| 2576 | pET22b-Δ25-*cwlD*$_{E199A}$-$His_6$ | pET22b-*cwlD* (deletion of N-terminal 25 aa and E199A point mutation with His6 tag) in BL21(DE3) | This study |
| 2587 | pET22b-Δ25-*cwlD*-$His_6$ + pET29a Δ22-*gerS*+TAA | Co-expression of pET22b-*cwlD* (deletion of N-terminal 25 aa with His6 tag) with pET29a-*gerS* (untagged deletion of N-terminal 22 aa) in BL21(DE3) | (3) |
| 2588 | pET22b-Δ25-*cwlD*$_{H49A}$-$His_6$ + pET29a Δ22-*gerS*+TAA | Co-expression of pET22b-*cwlD* (deletion of N-terminal 25 aa and H49A point mutation with His6 tag) with pET29a-*gerS* (untagged deletion of N-terminal 22 aa) in BL21(DE3) | This study |
| 2589 | pET22b-Δ25-*cwlD*$_{E65A}$-$His_6$ + pET29a Δ22-*gerS*+TAA | Co-expression of pET22b-*cwlD* (deletion of N-terminal 25 aa and E65A point mutation with His6 tag) with pET29a-*gerS* (untagged deletion of N-terminal 22 aa) in BL21(DE3) | This study |
| 2590 | pET22b-Δ25-*cwlD*$_{E199A}$-$His_6$ + pET29a Δ22-*gerS*+TAA | Co-expression of pET22b-*cwlD* (deletion of N-terminal 25 aa and E199A point mutation with His6 tag) with pET29a-*gerS* (untagged deletion of N-terminal 22 aa) in BL21(DE3) | This study |
| 2593 | pET22b-Δ25-*cwlD*-$His_6$ + pET29a Δ22-*gerS*+TAA | Co-expression of pET22b-*cwlD* (deletion of N-terminal 25 aa with His6 tag) with pET29a-*gerS* (untagged deletion of N-terminal 22 aa) in B834 (DE3) | This study |
| 2984 | pET22b-Δ25-*cwlD*$_{E78Q}$-$His_6$ + pET29a Δ22-*gerS*+TAA | Co-expression of pET22b-*cwlD* (deletion of N-terminal 25 aa and E78Q point mutation with His6 tag) with pET29a-*gerS* (untagged deletion of N-terminal 22 aa) in BL21(DE3) | This study |
| 2987 | pET22b-Δ25-*cwlD*$_{R169D}$-$His_6$ + pET29a Δ22-*gerS*+TAA | Co-expression of pET22b-*cwlD* (deletion of N-terminal 25 aa and R169D point mutation with His6 tag) with pET29a-*gerS* (untagged deletion of N-terminal 22 aa) in BL21(DE3) | This study |
| 2993 | pET22b-Δ25-*cwlD*-$His_6$ + pET29a Δ22-*gerS*$_{D106R}$+TAA | Co-expression of pET22b-*cwlD* (deletion of N-terminal 25 aa with His6 tag) with pET29a-*gerS* (untagged deletion of N-terminal 22 aa with D106R point mutation) in BL21(DE3) | This study |
| 2994 | pET22b-Δ25-*cwlD*-$His_6$ + pET29a Δ22-*gerS*$_{H61A}$+TAA | Co-expression of pET22b-*cwlD* (deletion of N-terminal 25 aa with His6 tag) with pET29a-*gerS* (untagged deletion of N-terminal 22 aa with H61A point mutation) in BL21(DE3) | This study |
| 2996 | pET22b-Δ25-*cwlD*$_{R169D}$-$His_6$ + pET29a Δ22-*gerS*$_{D106R}$+TAA | Co-expression of pET22b-*cwlD* (deletion of N-terminal 25 aa and R169D point mutation with His6 tag) with pET29a-*gerS* (untagged deletion of N-terminal 22 aa with D106R point mutation) in BL21(DE3) | This study |
| 3008 | pMTL-YN1C-*cwlD*$_{R169Q}$-3xFLAG | pMTL-YN1C-*cwlD*$_{R169Q}$-3xFLAG in HB101/pRK24 | This study |
| 3009 | pMTL-YN1C-*cwlD*$_{R169D}$-3xFLAG | pMTL-YN1C-*cwlD*$_{R169D}$-3xFLAG in HB101/pRK24 | This study |

| | | | |
|---|---|---|---|
| 3010 | pMTL-YN1C-*gerS*$_{D106N}$-3xFLAG-*alr2* | pMTL-YN1C-*gerS*$_{D106N}$-3xFLAG-*alr2* in HB101/pRK24 | This study |
| 3011 | pMTL-YN1C-*gerS*$_{D106R}$-3xFLAG-*alr2* | pMTL-YN1C-*gerS*$_{D106R}$-3xFLAG-*alr2* in HB101/pRK24 | This study |

### Plasmids

| | | |
|---|---|---|
| pET22b | For cloning His-tagged expression constructs | Novagen |
| pET29a | For cloning His-tagged expression constructs | Novagen |
| pMTL-YN1C | For cloning complementation constructs to be integrated into the pyrE locus of 630Δ*erm*Δ*pyrE* | (1) |

### Supplementary Table S2: Primers used in this study.

| Number | Primer Name | Primer Sequence |
|---|---|---|
| 2181 | 5' NotI *gerS* Gibson | GGAATTAGGGATGTAATAA**GCGGCCGC**CAGTTGTAGATTCAGAGAATAGAGTTG |
| 2362 | 5' NotI P$_{cwlD}$ Gibson | GGAATTAGGGATGTAATAA**GCGGCCGC**ATAAGGTTATATATTTTAAGATATTATTTAAC |
| 2365 | 3' gerS rev eos | CTCTAAAACCAAATGTCCTTTGGATAACTTTACTTTCATCTCCTC |
| 2408 | 3' XhoI *gerS* downstream Gibson | CAAGCTTGCATGTCTGCAGGC**CTCGAG**TTATTTTAGCAAATAACTGTTTATTTG |
| 2450 | 3' XhoI *cwlD* YN1C | CAAGCTTGCATGTCTGCAGGC**CTCGAG**TCAACTTAAATATTTTTGTATTCCTATGTAG |
| 2545 | 5' Δ25aa-*cwlD* Gibson NdeI pET22b | GTTTAACTTTAAGAAGGAGATATA**CATATG**AAAAATATTTCTGAAGATGTTATCAAG |
| 2546 | 3' Δ25aa-*cwlD* Gibson SalI pET22b | GCTCGAGTGCGGCCGCAAGCTT**GTCGAC**ACTTAAATATTTTTGTATTCCTATGTAGATTG |
| 2562 | 3' XhoI FLAG YN1C | CAAGCTTGCATGTCTGCAGGC**CTCGAG**TTATTTGTCATCATCATCCTTATAATCC |
| 2599 | 3' *cwlD*-3XFLAG rev eos | CATCCCTTGGTTTGACTTCTCTGTTGTTTGTCTTATCTACTACTCTTTTAAGTTCTTC |
| 2617 | 5' *gerS*-3xFLAG-alr2 Gibson SOE | GGATTATAAGGATGACGATGACAAATAAGGGGGACTAAAGACATGCAAAAAATAACAGTG |
| 2762 | 5' *cwlD* (H49A) SOE | CAAAACTATAATTTTAGATGCTGGTGCAGGAGGCATTGATCCAGGTGCATTAAATAAG |
| 2763 | 3' *cwlD* (H49A) eos | CTTATTTAATGCACCTGGATCAATGCCTCCTGCACCAGCATCTAAAATTATAGTTTTG |
| 3103 | Δ22-*gerS* Gibson NdeI pET22 | GTTTAACTTTAAGAAGGAGATATA**CATATG**CAAAAACGACAGTCCACAAAAGAAGAAG |
| 3104 | Δ22-*gerS* (no stop) Gibson SalI pet22b rev | CTCGAGTGCGGCCGCAAGCTT**GTCGAC**GTTTCTGTATTCAAAATCTTTGTATTTTAC |
| 3251 | 3' *cwlD* E199A eos | CACTCTTTTTCATTTGATAAAAAACCACATGCTATCAGTACTGATGGAATATTATT |
| 3252 | 5' c*wlD* E199A SOE | AATAATATTCCATCAGTACTGATAGCATGTGGTTTTTTATCAAATGAAAAAGAGTG |
| 3309 | 3' *cwlD* E65A eos | TAAGTGTTATTGCTAGGTTAATATCCTTTGCAGATGTACTCTTATCCTTATTTAATG |
| 3310 | 5' *cwlD* E65A SOE | CATTAAATAAGGATAAGAGTACATCTGCAAAGGATATTAACCTAGCAATAACACTTA |
| 3419 | 5’ Δ22aa-*gerS* NdeI pet29 | GTTTAACTTTAAGAAGGAGATATA**CATATG**AGAAAAAAGTGGACCATAGTATGTAT |
| 3420 | 3’ Δ22aa-*gerS* XhoI pet29 | GATCTCAGTGGTGGTGGTGGTGGTG**CTCGAG**TCAGTTTCTGTATTCAAAATCTTTGTAT |

| | | |
|---|---|---|
| 3719 | 3' gerS D106N | ATTTTTTCCTGTATTAGGTAACTCCACTACATTACTAATCTTTGGATT TTTAACTAATA |
| 3720 | 5' gerS D106N | TATTAGTTAAAAATCCAAAGATTAGTAATGTAGTGGAGTTACCTAAT ACAGGAAAAAAT |
| 3723 | 3' *gerS* D160R eos | TTTTTTCCTGTATTAGGTAACTCCACTACACGACTAATCTTTGGATTT TTAACTAATA |
| 3724 | 5' gerS D160R SOE | TATTAGTTAAAAATCCAAAGATTAGTCGTGTAGTGGAGTTACCTAAT ACAGGAAAAAA |
| 3725 | 3' *gerS* H61A eos | TTTTTTATAAGTATGAATTAAAACATAATTAGCTGGGCTTTTATTTCC AACTACTTCTA |
| 3726 | 5' *gerS* H61A SOE | TAGAAGTAGTTGGAAATAAAAGCCCAGCTAATTATGTTTTAATTCAT ACTTATAAAAAA |
| 3727 | 3' *cwlD* R169Q eos | GACTTCTCTGTTGTTTGTCTTATCTACTACTTGTTTAAGTTCTTCTTGA ATACACTTCG |
| 3728 | 5' *cwlD* R169Q SOE | CGAAGTGTATTCAAGAAGAACTTAAACAAGTAGTAGATAAGACAAA CAACAGAGAAGTC |
| 3731 | 3' *cwlD* R169D eos | GACTTCTCTGTTGTTTGTCTTATCTACTACATCTTTAAGTTCTTCTTGA ATACACTTCG |
| 3732 | 5' *cwlD* R169D SOE | CGAAGTGTATTCAAGAAGAACTTAAAGATGTAGTAGATAAGACAAA CAACAGAGAAGTC |
| 3733 | 3' *cwlD* E78Q eos | TATTACAAGGCCACCACTTGATTCTATAAGTTGTCTAAGCTTAAGTG TTATTGCTAGGT |
| 3734 | 5' *cwlD* E78Q SOE | ACCTAGCAATAACACTTAAGCTTAGACAACTTATAGAATCAAGTGGT GGCCTTGTAATA |

Restriction sites are underlined and in bold.

### Supplementary Table S3: Crystallographic data collection and structure refinement statistics for CwlD:GerS.

| | Native (*L8*) | SeMet (*R9*) | Iod (*R13*) |
|---|---|---|---|
| PDB ID | | *ref113* | |
| Data Processing Proteum | | | |
| Wavelength | 1.033 | 0.9794 | 0.9794 |
| Space Group<br>Cell a=b, c (Å) | P6(5)22<br>104.6, 156.2 | P6(5)22<br>105.2, 154.8 | P6(5)22<br>10.65, 156.5 |
| Resolution (Å)<br>(High resolution)[a] | 45 - 2.40<br>(2.48 - 2.40) | 45 - 2.40<br>(2.49-2.40) | 46 - 2.49<br>(2.58 - 2.59) |
| Reflections | 20346 | 20401 | 18805 |
| Completeness (%) | 99.5 (98.4) | 99.9 (99.2) | 99.5 (95.3) |
| Wilson B | 72.9 | 59.9 | 69.8 |
| Multiplicity | 19.0 (19,.4) | 18.7 (18.3) | 19.0 (19.6) |
| I/sigI | 14.1 (1.0) | 14.5 (1.5) | 14.4 (1.5) |
| $R_{meas}$ % | 15.1 (261) | 12.4 (231) | 10.3 (256) |
| $R_{pim}$ % | 3.5 (59.,1) | 2.9 (53.6) | 2.4 (57.6) |
| $CC^{1/2}$ | 0.997 (0.345) | 0.999 (0.572) | 0.999 (0.644) |
| Phasing | | | |
| Heavy Atom Sites Sharp45<br>$CC_{all}/CC_{weak}$<br>CFOM<br>Correlation | | 6<br>56.8/34.8<br>91.6<br>0.501 | 12<br>41.5/9.4<br>50.9<br>0.363 |

| | | | |
|---|---|---|---|
| Phasing Power<br>Isomorphous<br>Anomalous | | <br>1.02<br>2.73 | <br>0.64<br>0.64 |
| Phasing R-cullis acentric<br>Isomorphous<br>Anomalous | | <br>0.275<br>0.464 | <br>0.621<br>0.893 |
| Density Modification FOM | | 0.63 | |
| Model refinement | | | |
| $R_{work}/R_{free}$ % | | 20.3/24.9 | |
| RMS Bonds (Å)<br>RMS Angles (°) | | 0.008<br>1.00 | |
| Ramachandran (%)<br>Favored<br>Outlier | | <br>95.8<br>0.3 | |
| Mean B (Å$^2$) | | 69.2 | |
| Correlation Coefficient | | 0.823 | |
| Atoms Protein<br>Ligand ($Zn^{2+}$)<br>Solvent | | 2890<br>1<br>18 | |
| Coordinate error (Å)<br>Phase error (°) | | 0.39<br>30.1 | |

[a]Numbers in parentheses denote high resolution bin.
